# Studies of NH_4_^+^ and NO_3_^-^ uptake ability of subalpine plants and resource-use strategy identified by their functional traits

**DOI:** 10.1101/372235

**Authors:** Legay Nicolas, Grassein Fabrice, Arnoldi Cindy, Segura Raphaël, Laîné Philippe, Lavorel Sandra, Clément Jean-Christophe

## Abstract

The leaf economics spectrum (LES) is based on a suite of leaf traits related to plant functioning and ranges from resource-conservative to resource-acquisitive strategies. However, the relationships with root traits, and the associated belowground plant functioning such as N uptake, including nitrate (NO_3_^-^) and ammonium (NH_4_^+^), is still poorly known. Additionally, environmental variations occurring both in time and in space could uncouple LES from root traits. We explored, in subalpine grasslands, the relationships between leaf and root morphological traits for 3 dominant perennial grass species, and to what extent they contribute to the whole-plant economics spectrum. We also investigated the link between this spectrum and NO_3_^-^ and NH_4_^+^ uptake rates, as well as the variations of uptake across four grasslands differing by the land-use history at peak biomass and in autumn. Although poorly correlated with leaf traits, root traits contributed to an economic spectrum at the whole plant level. Higher NH_4_^+^ and NO_3_^-^ uptake abilities were associated with the resource-acquisitive strategy.

Nonetheless, NH_4_^+^ and NO_3_^-^ uptake within species varied between land-uses and with sampling time, suggesting that LES and plant traits are good, but still incomplete, descriptors of plant functioning. Although the NH_4_^+^: NO_3_^-^ uptake ratio was different between plant species in our study, they all showed a preference for NH_4_^+^, and particularly the most conservative species. Soil environmental variations between grasslands and sampling times may also drive to some extent the NH_4_^+^ and NO_3_^-^ uptake ability of species. Our results support the current efforts to build a more general framework including above- and below-ground processes when studying plant community functioning.

## Introduction

Functional traits have been widely used to describe different plant strategies. One major axis of specialisation involves a trade-off between conservation of resources in well protected and long lived tissues, and acquisition of resources by tissue with high use-efficiency and turn-over, and commonly referred as the leaf economic spectrum (LES, Wright *et al*. 2004). More specifically, species with an exploitative strategy share similar leaf attributes such as high specific leaf area (SLA) and nitrogen concentrations (LNC) that have been associated with short leaf life-span, high photosynthetic capacity as well as high decomposability (Reich 2014, Cornwell *et al*. 2008), and dominate in nutrient rich environments, while slow-growing conservative species carry opposite trait values and are more common in poor or harsh conditions (Chapin 1980, Ordonez et al. 2009). Despite some evidences of a similar contribution of root traits to the plant strategy (Roumet et al. 2006, Freschet et al. 2010, Fort et al. 2013), the significance of root traits is less understood than the one for leaf traits, mainly because weak correlations between analogous leaf and root traits have been reported (Craine et al 2005, Tjoelker et al. 2005, Freschet et al. 2010), and also because root functioning is often overlooked compared to leaves in field conditions. Nutrient uptake ability, one of the main functions provided by roots (Hodge 2004, James et al. 2009), is both influenced by anatomical and physiological adjustments such as specific root length or maximal uptake rate (Vmax, but see Bassirirad 2000). Among nutrients, nitrogen is one of the best studied mineral nutrients and its uptake by plants under both the ammonium (NH_4_^+^) and nitrate (NO_3_^-^) forms is influential for plant and ecosystem functioning. However, rarely have morphological and physiological properties of root been assessed simultaneously in field conditions, whereas some information come from species grown in standardized conditions (Maire et al. 2009, Grassein et al. 2015). NH_4_^+^ and NO_3_^-^ uptake can indeed be influenced by several environmental factors justifying the use of controlled conditions to estimate uptake parameters in a comparative purpose. For example, NH_4_^+^ and NO_3_^-^ uptake has been reported to vary in response to temperature or pH (Garnett and Smethurst 1999). Nevertheless, NH_4_^+^ and NO_3_^-^ uptake ability also differs between species, and is partially related to plant strategy and their functional traits (Grassein *et al*. 2015), but these results need to be validated for plant grown in natural conditions. Finally, NH_4_^+^ and NO_3_^-^ transporters have two components: a constitutive component and a component induced by the presence of NH_4_^+^ and NO_3_^-^in the soil solution. Thus, it is important to study interspecific differences for NH_4_^+^ and NO_3_^-^ uptake at a given site. Otherwise, it is difficult to interpret differences as resulting from species differences.

Subalpine grasslands are subject to the combined effects of climate and anthropogenic factors, both influencing N cycling and thus N availability for organisms (Bardgett *et al*. 2005, Legay *et al*. 2013). Decreased management intensity favours plant species with resource conservative traits (Quétier *et al*. 2007), which are usually associated with fungal-dominated belowground communities (de Vries *et al*. 2012, Grigulis *et al*. 2013). Concomitantly, it slows down N cycling (Zeller *et al*.2000, Robson *et al*. 2010), favouring the accumulation of soil ammonium (NH_4_^+^) rather than soil nitrate (NO_3_^-^) (Robson *et al*. 2007). Plants growing in such variable conditions are likely to adjust their N uptake ability, as it has been shown for functional traits (Quétier *et al*. 2007, Grassein et al. 2015).

In this study, we investigated the relationships between functional traits and inorganic N (NH_4_^+^ and NO_3_^-^) uptake for three perennial grass species with contrasted leaf economic strategies. Because soil inorganic NH_4_^+^ and NO_3_^-^ availability and plant NH_4_^+^ and NO_3_^-^ uptake ability are likely to vary across seasons and in response to management (Jaeger *et al*. 1999, Miller *et al*. 2009), we examined these relationships for individuals occurring in four subalpine grasslands with different management and throughout the growing season, thereby testing their temporal consistency. Estimating root NH_4_^+^ and NO_3_^-^ uptake, and measuring functional traits for leaves and roots, we tested the following hypotheses: (1) similar to leaf traits, root traits are also contributing to the plant economics spectrum with root traits reflecting nutrient acquisition (e.g. high specific root length and root nitrogen content) expected to be more associated to the exploitative syndrome, (2) and with more exploitative species being more efficient to take up both NH_4_^+^ and NO_3_^-^. (3) As functional traits are influenced by environmental conditions, we hypothesised that NH_4_^+^ and NO_3_^-^ uptake will be influenced by environmental variations between grasslands, as well as during the growing season, probably following NH_4_^+^ and NO_3_^-^ availability depending on the most abundant form.

## Methods

### Study site and species

The study site is located in the upper Romanche valley of the central French Alps between the village of Villar d’Arêne and the Lautaret Pass (Table 1). The climate is subalpine with a strong continental influence. Winters are cold and snowy, with monthly average minimum temperatures of −15.9°C in February, maximum monthly average temperature of 23.8°C in July, and mean annual precipitation of 956mm (unpublished data, sajf.ujf-grenoble.fr). The growing season starts following snow melt in late April - early May and continues until late September or October depending on the date of the first snow in autumn.

**Table 1.**
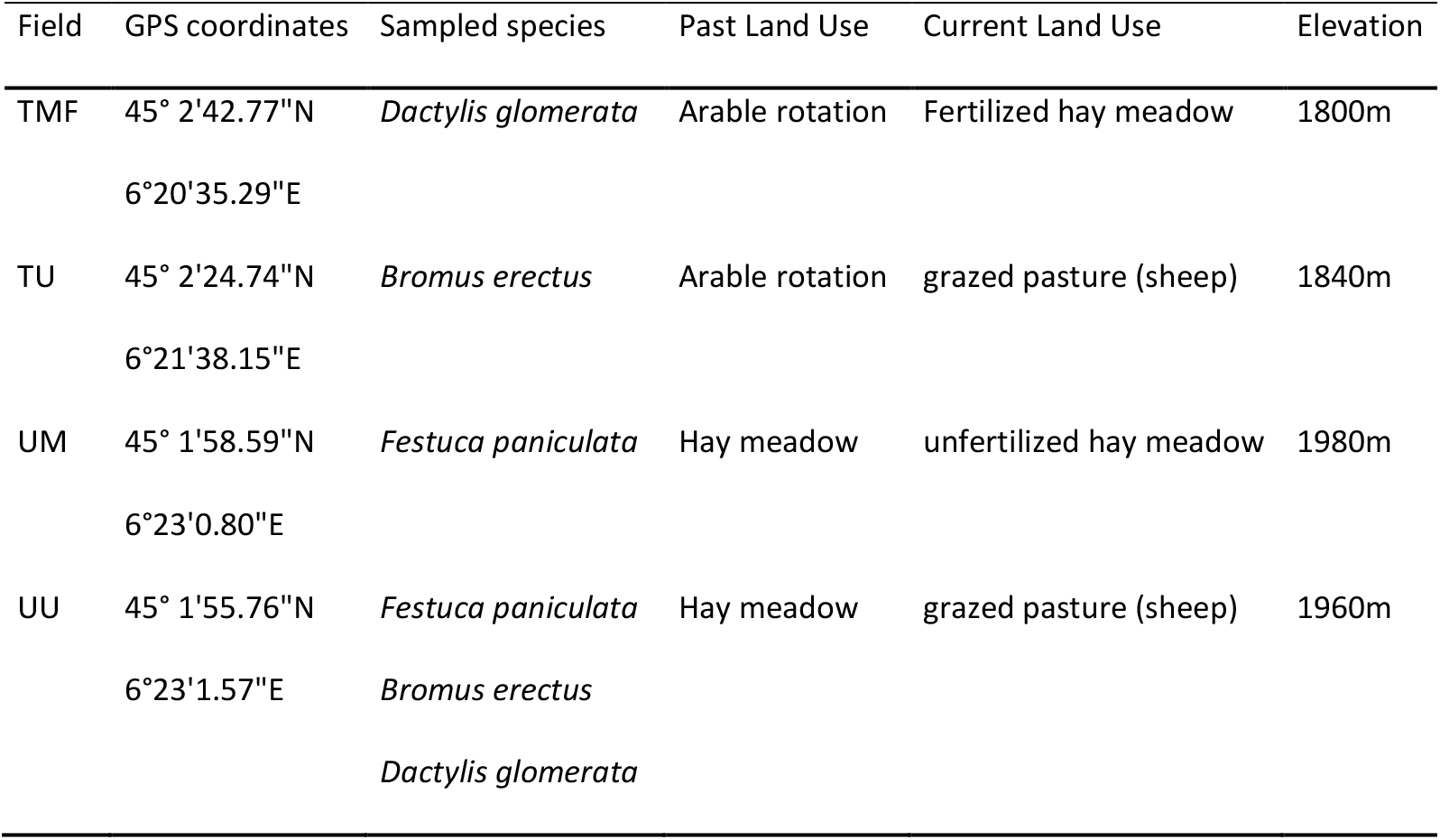
Description of the studied grasslands. Past and current land uses describe the former and current management of these grasslands (see Quetier et al. 2006 for more information). TMF: Terraced Mown and Fertilized, TU: Terraced Unmown not fertilized but lightly grazed, TM: Unterraced Mown, UU: Unterraced Unmown but lightly grazed.

Given the hypothesis that NH_4_^+^ and NO_3_^-^ uptake could be an important hard plant trait related to resource use strategy (as suggested by soft structural and morphological traits) and to field dominance, and due to the degree of precision chosen for NH_4_^+^ and NO_3_^-^ uptake estimations (see 2.2), a compromise was necessary regarding the number of species, grasslands and replicates to be investigated. This sampling adjustment was required to conduct N uptake estimations for all individuals in a brief enough time period so that most abiotic factors remained as comparable as possible (soil moisture, temperature, radiation).

We chose three common and dominant grass species, *Dactylis glomerata* L., *Bromopsis erecta* (Huds.) Fourr. (formerly *Bromus erectus* (Huds.)) and *Patzkea paniculata* (L.) G.H.Loos (formerly *Festuca paniculata* (L.) Schinz & Thell.). All species are perennial, arbuscular mycorrhizal non-dependent species and span a gradient from more exploitative (*D. glomerata*) to more conservative (*F. paniculata*) resource use strategies (Grassein et al. 2015). Four grasslands (Table 1), described in Quétier *et al*. (2007), were chosen for their contrasting past and current managements, and were similar to the grasslands studied by Robson *et al*. (2007, 2010): (i) Terraced Mown and Fertilized (TMF), (ii) Terraced Unmown not fertilized but lightly grazed (TU), (iii) Un-terraced Mown grassland (UM) and (iv) Un-terraced Unmown but lightly grazed grassland (UU), representing a gradient of decreasing management intensity. To reflect field dominance patterns, *D. glomerata* was sampled in TMF, *B. erectus* in TU, *F. paniculata* in UM, and all three species were sampled in UU where they coexist, although *F. paniculata* was dominant (Table 1).

To assess NH_4_^+^ and NO_3_^-^ uptake patterns over the growing season, the same sampling design was repeated twice during 2010. At each date for each species and grassland, we sampled the roots and soil (approximately: 25×25×25 cm) of five individuals (genetically distinct individuals at least 2m apart). The first sampling corresponded to the peak biomass and targeted flowering onset (just before anthesis), and the second sampling corresponded to autumn after last management activities occurred. For *D. glomerata* in TMF and *B. erectus* in TU, the two sampling dates were mid-June and mid-September. For *F. paniculata* in UM and the three species in UU, the sampling dates were: early July and early September. These two dates are called “Summer” and “Autumn” hereafter. As much as possible, species were sampled at the same time during the day to avoid any diurnal variation in N uptake (Gessler et al. 1998). In total, we have sampled 12 points (3 species*2 seasons*2 habitats per species).

### Soil nitrogen pools

At each date and for each grassland, soil nitrogen concentrations were measured from six soil cores (dimensions 4.5 cm Ø, 10 cm deep) kept on ice in the field and maintained at 4°C upon return to the laboratory (within 2h). Soils were sieved through a 5.6 mm mesh to remove roots and stones. A subsample of 10g fresh sieved soil was prepared for extraction of inorganic N in 0.5M K_2_SO_4_, and analysed using a colorimetric analyser (FS-IV autoanalyser (OI-Analytical, College Station, TX, USA) (following Bowman *et al*. 2003) to measure soil concentrations of ammonium (NH_4_^+^), nitrate (NO_3_^-^) and Total Dissolved Nitrogen (TDN). Soil aliquots were used to determine soil water (7 days at 70°C) and soil organic matter contents (550°C during 4 hours). Finally, soil subsamples were air-dried to measure soil pH, or ground to a fine powder for measurements of total carbon (C) and N contents using an elemental analyser (FlashEA 1112, Thermo Fisher Scientific Inc., Waltham, MA, USA).

At each date, five individuals of each species, with roots and soil, were excavated from each field, transferred within half an hour to the laboratory located at the Lautaret Pass (Station Alpine Joseph Fourier) and kept at 4°C until the NO_3_^-^ and NH_4_^+^ uptake rate measurements to maintain the functional integrity of the roots. Living young fine roots were washed with deionised water, cut to 2-cm length and then, rinsed in 1mM CaSO_4_ at 4°C for 3 min. The NO_3_^-^ and NH_4_^+^ uptake rates were measured during the first hour following plant harvest as described by Louahlia *et al*. (2000). The optimal conditions for uptake measurements by excised root determined by Lainé *et al*. (1993) were used in the present study.

### Functional traits

Functional traits were measured for roots and leaves using standardised protocols (Perez-Harguindeguy *et al*. 2013). Two of the individual root sub-samples were used to estimate root dry matter content (RDMC), specific root length (SRL, Winrhizo^®^ software, fresh length per unit of dry mass), and were further analysed to obtain root ^15^N natural abundance and root nitrogen concentration (RNC, N mass per unit of dry mass). Specific leaf area (SLA, fresh area per unit of dry mass), leaf and root dry matter contents (LDMC and RDMC, dry mass per unit of fresh mass), leaf nitrogen concentration (LNC, N mass per unit of dry mass) were also measured.

### Nitrogen uptake estimation: the “excised” roots method

Although measuring only a net N uptake, which is the result of influx and efflux, the direct measurement of N uptake using excised roots allows characterising the plant uptake kinetics for NO_3_^-^ and NH_4_^+^ while controlling for the environmental variations. This makes it possible to compare different species at the cost of losing relevant ecological information (Lucash *et al*. 2007). This method was thus applied to plants collected in the field. Root N uptake kinetics started within 60 min after excision, thereby avoiding the potential decline in N uptake ability reported to start after 3h (Louahlia *et al*. 2000). Nitrate and ammonium uptake by plants involved mainly the transport system called HATS (High Affinity Transport System). It contributes to N uptake at low to moderate concentrations of external N (<1mM) and saturates at 0.2-0.5 mM (Kronzucker *et al*. 1999, Min *et al*. 2000), which makes it the more likely system used by plants growing in natural and semi-natural ecosystems limited by N (Bassirirad 2000, Maire *et al*. 2009). The estimation of the maximum NH_4_^+^ and NO_3_^-^ uptake rates by HATS requires a range of N concentrations below 1mM at which the Vmax can be reached depending on species (Grassein *et al*. 2015). Consequently, uptake was estimated from the accumulation of ^15^N in root sub-samples incubated for one hour in a buffer solution (pH = 5.5-following Leon *et al*. (1995)), containing a range of N concentrations (20, 50, 100, 250, 500 and 1000 μM). Six sub-samples were incubated in K^15^NO_3_ and the other six in (^15^NH_4_)_2_SO_4_ with a ^15^N excess of 99% atom. The two N forms were tested individually in order to avoid possible interactions (Kronzucker *et al*. 1999). Solution volumes and fresh weights were selected to avoid N depletion during the experiment. After 1h incubation, roots were washed twice for one minute with a 1mM CaSO4 at 4°C to stop any metabolic processes. Roots were then dried at 60°C for 72h, ground to a fine powder and analysed by IRMS at the University of Caen (Isoprime GV instruments, Stockport, UK) to obtain ^15^N Atom% and N concentrations.

### Data analysis

Nitrogen Uptake Rate (NUR) was calculated for each concentration and each inorganic N form (NH_4_^+^ and NO_3_^-^) using the ^15^N increase in the root incubated compared to the non-incubated control, and expressed by unit of time and dry mass (nmolN.h^-1^.g^-1^ of dry roots, see Leon *et al*. 1995). The dependence of NUR on substrate concentration was fitted for each individual and Hanes’s relation (Michaelis transformation) was used to estimate the maximum uptake rate (Vmax) defined as the maximum NUR for NH_4_^+^ and NO_3_^-^(Leon *et al*. 1995). Finally, the NH_4_^+^:NO_3_^-^ uptake ratio was calculated as the ratio between NH_4_^+^ Vmax and NO_3_^-^ Vmax.

A principal component analysis (PCA) was performed using all plant functional traits at the individual level to describe their functional strategy based on leaf and root traits. To investigate the relationships between functional traits of leaves and roots, and NH_4_^+^ and NO_3_^-^ uptake ability (hypothesis 1), we used Pearson correlation coefficients. Relationships between the functional strategy and uptake of NH_4_^+^ and NO_3_^-^at the root level were tested using regression analyses between the N uptake rates (Vmax) and the first PCA.

Comparisons of NH_4_^+^ and NO_3_^-^ uptake rates for species (hypothesis 2), fields and date (hypothesis 3) were conducted with ANOVA followed by Tukey tests to compare species and grasslands. In details, the effects of sampling time and fields on plant traits within each species were tested using two-ways ANOVA. Similarly, the effects of sampling time and fields on maximal NH_4_^+^ and NO_3_^-^ uptake rates within each species were tested using two-ways ANOVA. The effects of fields and sampling time on NH_4_^+^:NO_3_^-^ ratio within each species were tested using two-ways ANOVA. Then, we tested only in UU grasslands, the species effect using one-way ANOVA. Finally, we used a two-ways ANOVA and Tukey post hoc test to test soil parameters differences between fields and dates. Data were log-transformed when necessary to achieve normality and heteroscedasticity. All statistical analyses were performed using the software R 3.4.4, with multivariate analyses (PCA) being performed using the package Ade4 (Dray & Dufour 2007).

## Results

We observed large variations for leaf and root functional traits in spite of a restricted number of species in our study (Table 2). The range of variation was similar to, and sometimes even higher than the variability reported in Fort *et al*. (2013) for a larger set of species occurring in a similar ecosystem, including *D. glomerata* and *B. erectus*. The PCA of functional traits highlighted a first axis explaining 62.1% of the total variance (Fig. 1). The three species differed significantly for their mean position along this axis (p=0.012), with positive values for *D. glomerata* and negative values for *F. paniculata*. Positive values along this axis were characterised by high SLA, LNC and SRL, and low LDMC. Among these, SLA and LNC have been reported as major contributors to a resource economic spectrum establishing the existence of a fundamental trade-off between plant features allowing resource capture and those allowing resource conservation.

**Figure 1.**
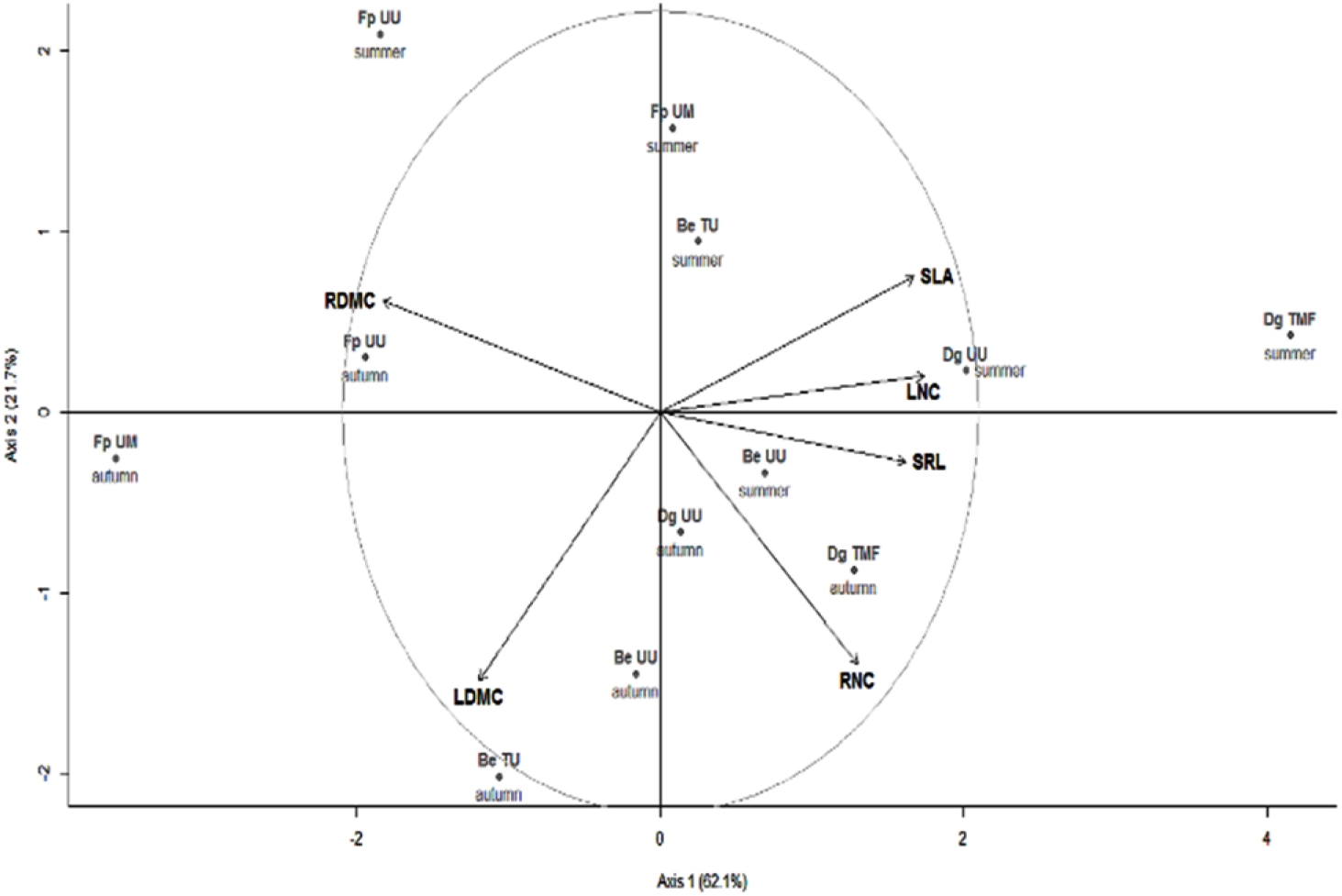
Principal components analysis (PCA) of functional traits measured for the leaves and roots of three grass species (Be: *Bromus erectus*, Dg: *Dactylis glomerata* and Fp: *Festuca paniculata*), in each grassland with different management (UU: unterraced unmown, UM: unterraced mown, TMF: terraced mown and fertilized, TU: terraced unmown). SLA: Specific leaf area, LDMC: Leaf dry matter content, LNC: Leaf nitrogen content, SRL: Specific root length, RDMC: Root dry matter content, RNC: Root nitrogen content.

**Table 2.** Mean values ± standard errors of leaf and root traits for each species, site and sampling time (n=5). For a given trait and species, statistically similar values have the same letter (Tukey post-hoc test). Bold values indicate the season with the highest trait values for a given species in a given grassland.

| Species | Site | Season | SLA ($\text{mm}^2.\text{g}^{-1}$) | LDMC ($\text{mg}.\text{g}^{-1}$) | LNC ($\text{mg}.\text{g}^{-1}$) | SRL ($\text{m}.\text{g}^{-1}$) | RDMC ($\text{mg}.\text{g}^{-1}$) | RNC ($\text{mg}.\text{g}^{-1}$) |
| --- | --- | --- | --- | --- | --- | --- | --- | --- |
| *B. erectus* | TU | Summer | **$30.07 \pm 0.8^a$** | $277.6 \pm 2.1^c$ | $21.46 \pm 0.2$ | $249.86 \pm 72.0$ | $307.77 \pm 22.4$ | $6.06 \pm 0.1$ |
| | | Autumn | $15.43 \pm 1.1^c$ | **$444.06 \pm 16.8^a$** | $12.09 \pm 0.9$ | $253.58 \pm 34.3$ | $303.5 \pm 6.3$ | $7.68 \pm 0.4$ |
| | UU | Summer | **$22.58 \pm 0.7^b$** | $305.06 \pm 4.2^c$ | $15.57 \pm 0.8$ | $328 \pm 74.2$ | $290.82 \pm 21.7$ | $7.17 \pm 0.1$ |
| | | Autumn | $16.37 \pm 1^c$ | **$385.1 \pm 11.5^b$** | $19.28 \pm 2.7$ | $222.12 \pm 75.4$ | $288.32 \pm 21.6$ | $7.87 \pm 0.5$ |
| *D. glomerata* | TMF | Summer | **$34.06 \pm 1.6^a$** | $248.79 \pm 17.7^c$ | **$32.6 \pm 3.4^a$** | $426.22 \pm 60.5$ | $232.49 \pm 20.5^b$ | $7.28 \pm 0.4$ |
| | | Autumn | $20.15 \pm 2^b$ | **$298.51 \pm 2.5^b$** | $19.41 \pm 2.2^b$ | $290 \pm 48.6$ | $249.09 \pm 8^b$ | $7.99 \pm 0.3$ |
| | UU | Summer | $26.17 \pm 0.3^{ab}$ | $265.78 \pm 9.1^{bc}$ | **$24.2 \pm 1.4^{ab}$** | $318.69 \pm 63.8$ | $254.08 \pm 5.6^b$ | $6.91 \pm 0.5$ |
| | | Autumn | $24.34 \pm 1.7^b$ | **$389.69 \pm 10.5^a$** | $21.81 \pm 2.5^{ab}$ | $216.87 \pm 44.7$ | **$304.99 \pm 8.8^a$** | $7.16 \pm 0.2$ |
| *F. paniculata* | UM | Summer | **$23.26 \pm 0.4^a$** | $232.06 \pm 0.6^c$ | **$20.22 \pm 1.9^a$** | $195.85 \pm 21.3^{ab}$ | $300.1 \pm 21.5$ | $5.15 \pm 0.4$ |
| | | Autumn | $8.78 \pm 0.5^b$ | **$433.87 \pm 18.8^a$** | $10.61 \pm 1.4^b$ | $153.37 \pm 20.5^b$ | $369.12 \pm 21.7$ | $4.32 \pm 0.5$ |
| | UU | Summer | **$21.36 \pm 0.9^a$** | $261.06 \pm 15.6^c$ | $14.52 \pm 1.8^{ab}$ | $166.21 \pm 20.7^{ab}$ | $380.91 \pm 6.4$ | $4.16 \pm 0.1$ |
| | | Autumn | $11.76 \pm 1.4^b$ | **$369.71 \pm 7.2^b$** | $15.53 \pm 3.6^{ab}$ | $293.71 \pm 54.5^a$ | $366.84 \pm 23.3$ | $4.31 \pm 0.1$ |

This functional axis was positively correlated to NH_4_^+^ and NO_3_^-^ Vmax (Fig 2a and 2b) and negatively to NH_4_^+^:NO_3_^-^ uptake ratio (Fig 2c) indicating a more pronounced preference for NH_4_^+^ at lower values of axis 1. Except RDMC, all traits taken separately were poorer predictors of the NO_3_^-^ and NH_4_^+^ maximum uptake rates than this functional axis, although the first PCA axis was significantly correlated with all functional traits (Table 3).

**Figure 2.**
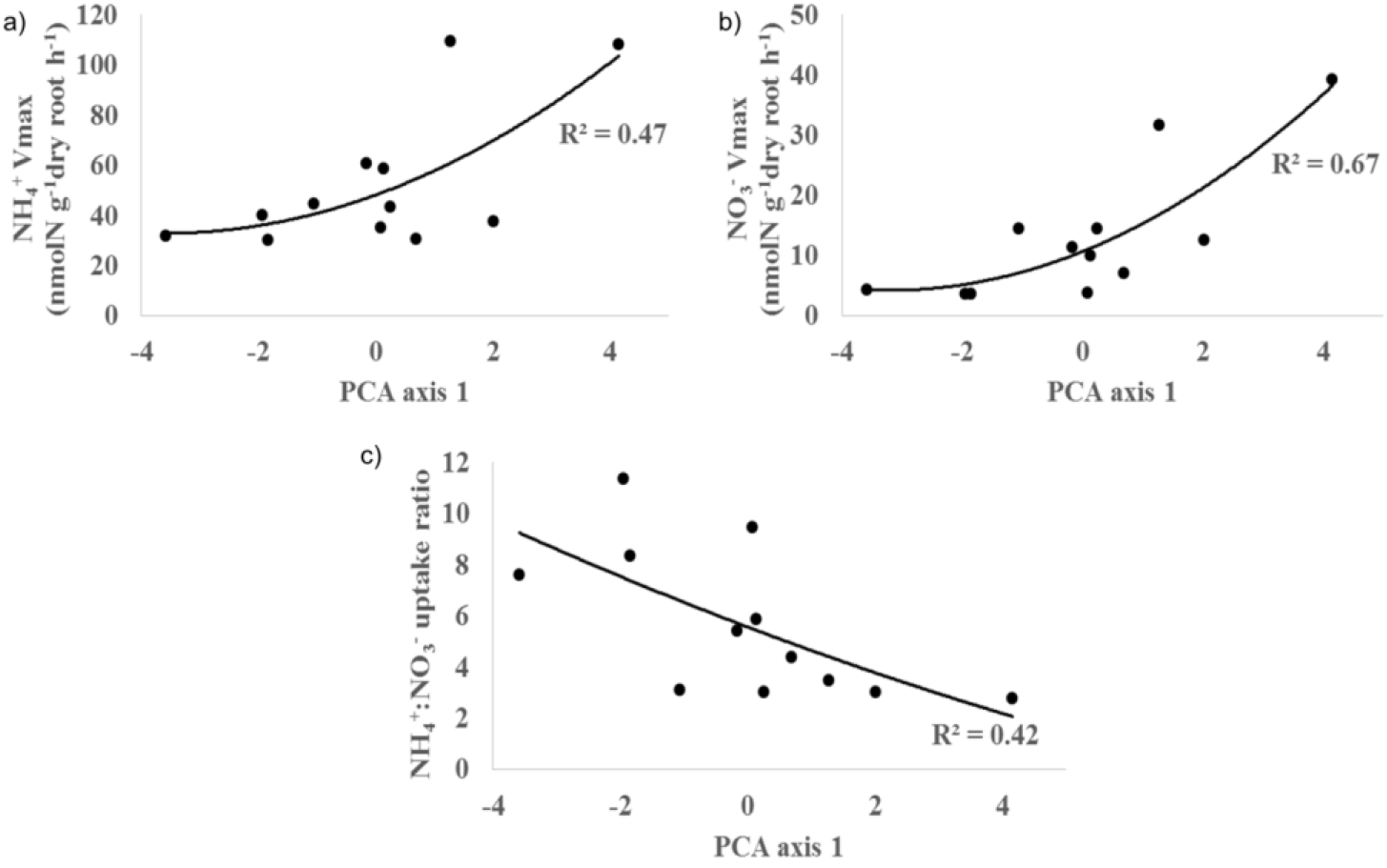
Relationships between the first axes of the PCA (fig 1) and Vmax for NH_4_^+^ (a), NO_3_^-^ (b) and NH_4_^+^:NO_3_^-^ uptake ratio (c). The three relationships were significant (p-values<0.05) assuming a polynomial relationship of order=2, and the resulting R^2^ are indicated on each graph.

**Table 3.** Pearson correlations between NH_4_^+^ and NO_3_^-^maximum uptake rates (Vmax), PCA axes, leaf and root traits. Significant values (p-value <0.05) are indicated in bold.

| | Vmax$\text{NH}_4^+$ | Vmax$\text{NO}_3^-$ | axe1 | axe2 | ratio | SLA | LDMC | SRL | RDMC | LNC |
| --- | --- | --- | --- | --- | --- | --- | --- | --- | --- | --- |
| Vmax$\text{NO}_3^-$ | **0.93** | | | | | | | | | |
| axe1 | **0.65** | **0.76** |  |  |  |  |  |  |  |  |
| axe2 | -0.28 | -0.24 | 0.00 |  |  |  |  |  |  |  |
| ratio | -0.46 | **-0.68** | **-0.64** | 0.43 |  |  |  |  |  |  |
| SLA | 0.37 | 0.53 | **0.84** | 0.38 | -0.54 |  |  |  |  |  |
| LDMC | -0.17 | -0.26 | **-0.60** | **-0.74** | 0.09 | **-0.74** |  |  |  |  |
| SRL | 0.55 | **0.70** | **0.81** | -0.14 | -0.51 | 0.54 | -0.34 |  |  |  |
| RDMC | **-0.68** | **-0.77** | **-0.91** | 0.31 | **0.75** | **-0.63** | 0.37 | **-0.71** |  |  |
| LNC | **0.64** | **0.65** | **0.87** | 0.10 | -0.32 | **0.69** | -0.51 | **0.66** | **-0.74** |  |
| RNC | **0.58** | **0.62** | **0.65** | **-0.70** | **-0.80** | 0.34 | 0.08 | 0.50 | **-0.83** | 0.42 |

In UU grassland, NH_4_^+^ Vmax in summer was similar for the three species (Fig. 3a) but greater for *D. glomerata* for NO_3_^-^ Vmax (p <0.001, Fig. 3b). Vmax in autumn for both N forms was lower for *F. paniculata* compared to the two other species (NH_4_^+^ p<0.05, NO_3_^-^ p< 0.001). Comparing the different grasslands within species, we observed reduced NO_3_^-^ and NH_4_^+^ Vmax values in the UU grassland for *D. glomerata* (in summer and in autumn) and *B. erectus* (summer) compared to the other grasslands. On the other hand, highest NH_4_^+^ Vmax for *B. erectus* and *F. paniculata* were found in UU during the autumn. Illustrating the seasonal variability, all species in the UU grassland had higher NH_4_^+^ maximum uptake rates in the autumn than in the summer, as well as higher NO_3_^-^ uptake for *B. erectus*. NH_4_^+^:NO_3_^-^ uptake ratio did not vary in time, but always showed higher values in the UU for the three species compared to the other grasslands, and overall greater values for *F. paniculata* (Fig. 4).

**Figure 3.**
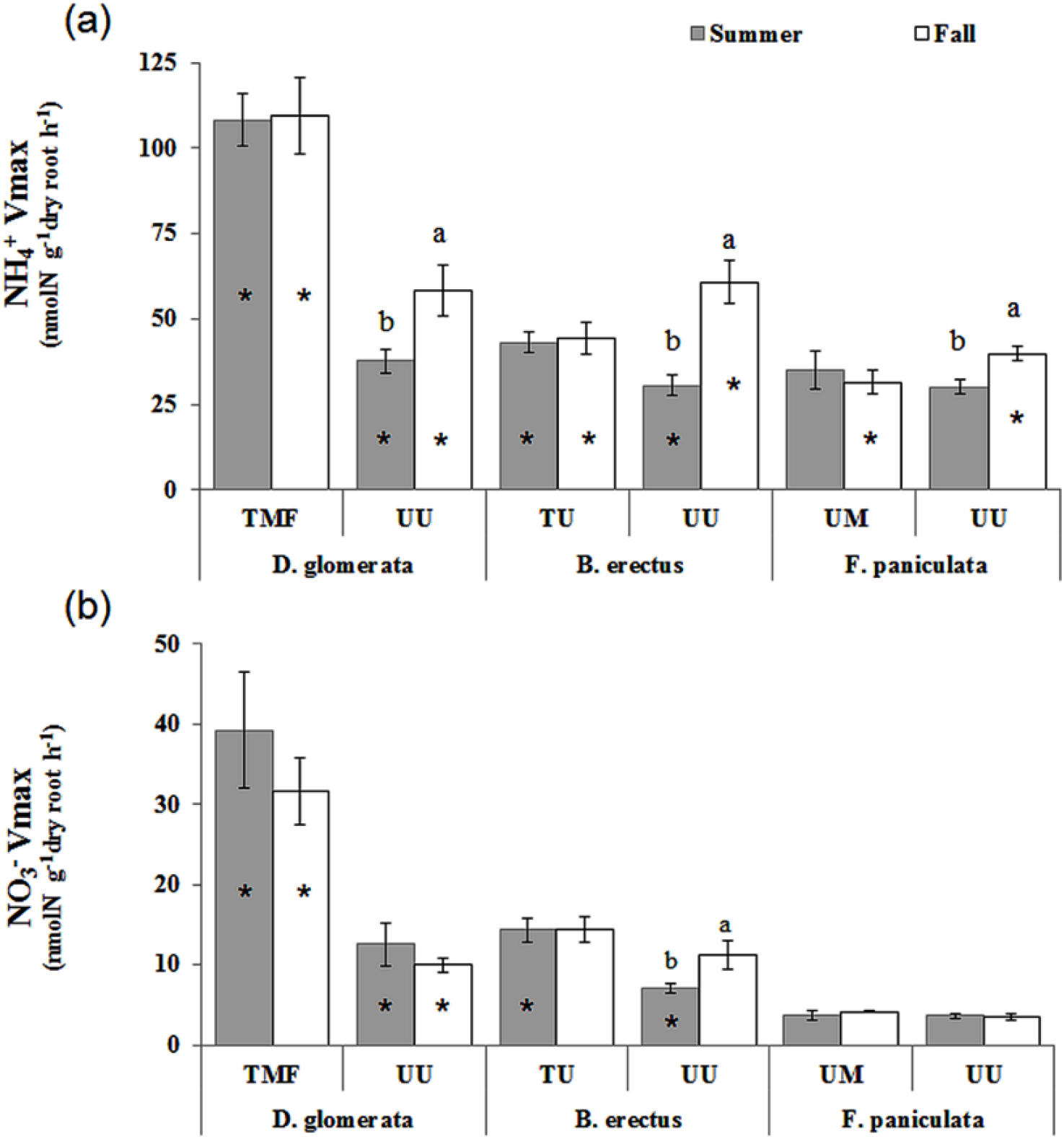
Vmax (Maximal uptake rate) for NH_4_^+^ (a) and NO_3_^-^ (b) of *D. glomerata, B. erectus* and *F. paniculata*. Within each combination of site and species, dates with the same letter had similar uptake parameters (Tukey post hoc test at 5%level, after an Anova with date as main effect). For each species, the significance of the differences between the two sites for uptake parameters were tested using a Student test, and stars indicate the dates at which the two sites differ significantly with a p-value<0.05.

**Figure 4.**
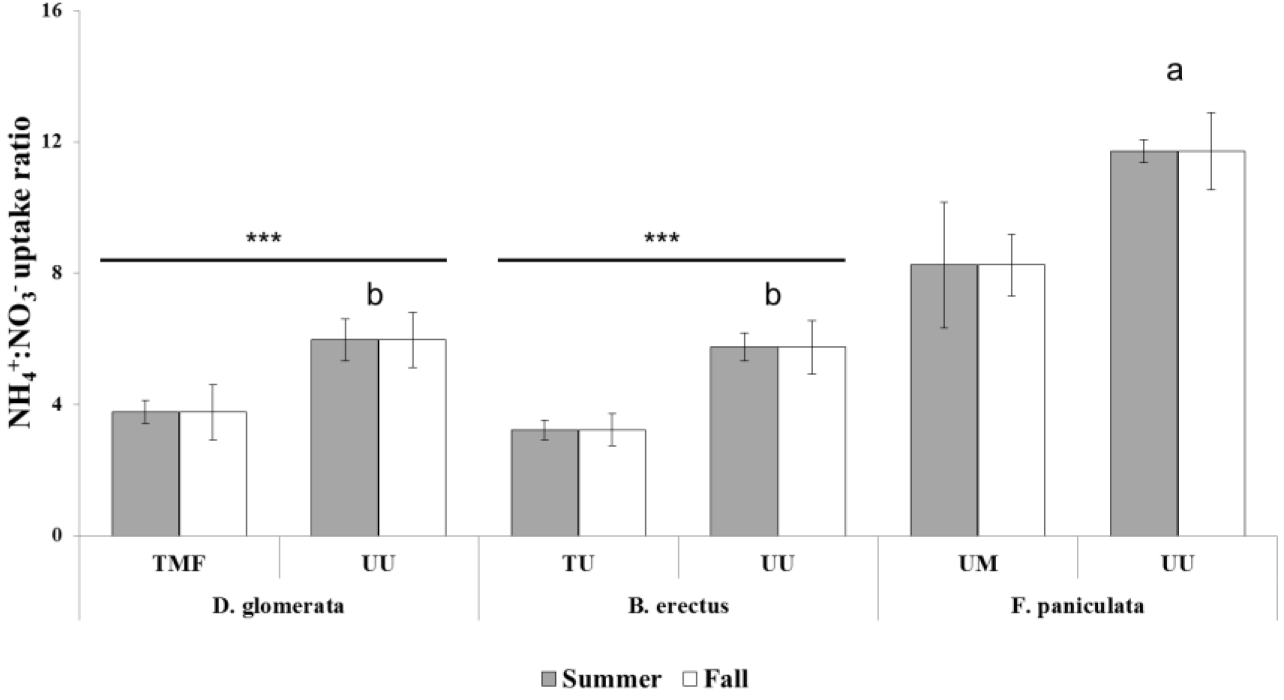
NH_4_^+^:NO_3_^-^ uptake ratio for the three species, in each site and at the two sampling times. The uptake ratio is unitless (ratio between NH_4_^+^ Vmax and NO_3_^-^Vmax). Within each species, *or *** indicate significant site effects (p value < 0.05 and 0.001 respectively) within each species (two-ways ANOVA with site, date and the interaction as main effect). In the grassland (UU) where all species occurred, the differences between species and sampling time were tested using two-ways ANOVA. Similar letters connect species with similar values in the UU grassland at both sampling dates.

Within species, a limited number of traits were significantly different between grasslands (Table 2). We only observed significant differences in autumn, with highest LDMC in TU for *B. erectus*, highest LDMC and RDMC in UU for *D. glomerata*, and highest LDMC but lowest SRL for *F. paniculata* in UM. However, changes in response to the season were more consistent among species and grasslands, with an increase of LDMC and a decrease of SLA in autumn compared to the summer. We also observed higher LNC for *D. glomerata* and *F. paniculata* during the summer than during the autumn in TMF and UM respectively, and higher RDMC during the autumn for *D. glomerata* in UU.

Since all species occurred in the UU grasslands, we choose to focus on soil parameters from these grasslands. UU and UM only differ for SWC in autumn (Table 4), all other soil variables were similar between these two grasslands, which had similar past land-use history (Table1). UU had consistently higher SWC and SOM, and lower soil pH and CN ratio than TMF and TU. All grasslands had similar soil NH_4_^+^ concentrations. During the summer, we observed higher TDN and NH_4_^+^:NO_3_^-^ soil ratio, and lower soil NO_3_^-^ concentration in UU compared to TMF and TU, but we did not find these differences in autumn.

**Table 4.** Soil properties (mean values ± SE) for each grassland and at each sampling time. No significant difference values between sites at a given date are shown by the same letter (Tukey post-hoc test). Values in bold indicate the highest values when the considered soil parameter was significantly different between dates in a grassland. na: not available because of a sampling issue. nd: not detectable: under the level of sensitivity of the method; TDN: total dissolved nitrogen).

|  |  | TMF | TU | UM | UU |
| --- | --- | --- | --- | --- | --- |
| Soil Water Content | Summer | **22.33 ± 0.71<sup>b</sup>** | **18.33 ± 1.12<sup>c</sup>** | na | **34.89 ± 1.20<sup>a</sup>** |
| (%) | Autumn | 11.44 ± 0.33<sup>b</sup> | 7.25 ± 0.88<sup>c</sup> | 13.58 ± 1.18<sup>b</sup> | 17.74 ± 1.54<sup>a</sup> |
| pH | Summer | 8.01 ± 0.04<sup>a</sup> | 8.03 ± 0.04<sup>a</sup> | n.a. | **6.31 ± 0.05<sup>b</sup>** |
| | Autumn | 7.98 ± 0.02<sup>a</sup> | 8.05 ± 0.04<sup>a</sup> | 5.85 ± 0.05<sup>b</sup> | 6.02 ± 0.08<sup>b</sup> |
| Soil Organic Matter | Summer | 13.16 ± 0.48<sup>c</sup> | 14.39 ± 0.93<sup>b</sup> | n.a. | 18.42 ± 0.64<sup>a</sup> |
| (%) | Autumn | 12.47 ± 0.65<sup>b</sup> | 11.38 ± 0.93<sup>b</sup> | 14.02 ± 0.43<sup>ab</sup> | 16.98 ± 1.29<sup>a</sup> |
| C:N ratio | Summer | **14.91 ± 0.35<sup>a</sup>** | 14.02 ± 0.64<sup>a</sup> | n.a. | 12.10 ± 0.15<sup>b</sup> |
| | Autumn | 13.46 ± 0.39<sup>a</sup> | 13.43 ± 0.64<sup>ab</sup> | 11.63 ± 0.14<sup>b</sup> | 11.74 ± 0.43<sup>b</sup> |
| TDN ($\mu\text{gN.g}^{-1}$ soil) | Summer | 20.83 ± 2.95<sup>b</sup> | 16.85 ± 1.33<sup>b</sup> | n.a. | 55.59 ± 12.15<sup>a</sup> |
| | Autumn | **46.12 ± 4.44<sup>a</sup>** | **46.13 ± 4.74<sup>a</sup>** | 34.54 ± 1.21<sup>ab</sup> | 46.65 ± 6.04<sup>a</sup> |
| NO<sub>3</sub><sup>-</sup> content | Summer | 3.87 ± 0.48<sup>ab</sup> | **5.63 ± 0.64<sup>a</sup>** | n.a. | 2.31 ± 0.53<sup>b</sup> |
| ($\mu\text{gN.g}^{-1}$ soil) | Autumn | 2.99 ± 0.54<sup>a</sup> | 0.54 ± 0.10<sup>b</sup> | n.d. | 1.39 ± 0.66<sup>ab</sup> |
| NH<sub>4</sub><sup>+</sup> content | Summer | 11.19 ± 0.47<sup>a</sup> | 15.43 ± 1.82<sup>a</sup> | n.a. | 12.23 ± 1.77<sup>a</sup> |
| ($\mu\text{gN.g}^{-1}$ soil) | Autumn | 10.07 ± 1.37<sup>a</sup> | 11.54 ± 1.55<sup>a</sup> | 6.97 ± 0.94<sup>a</sup> | 10.17 ± 1.92<sup>a</sup> |
| NH<sub>4</sub><sup>+</sup>:NO<sub>3</sub><sup>-</sup> ratio | Summer | 3.56 ± 0.65<sup>b</sup> | 2.97 ± 0.30<sup>b</sup> | n.a. | 7.85 ± 1.62<sup>a</sup> |
| | Autumn | 3.53 ± 0.36<sup>b</sup> | **28.68 ± 10.7<sup>a</sup>** | n.d. | 12.9 ± 6.4<sup>ab</sup> |

## Discussion

### Relationships between leaf and root traits

In the aim to find parallels between above and below-ground organs (e.g. Roumet et al. 2006), several studies have investigated the relationships between analogous traits measured for leaves and roots. While positive relationships have been reported for SLA *vs* SRL (Craine & Lee 2003, Craine et al. 2005, Freschet et al. 2010), other studies have reported a lack of relationships between SLA *vs* SRL (Craine et al. 2001, Tjoelker et al. 2005). In our study, we did not find any relationships between SRL/SLA, LDMC/RDMC and LNC/RNC, and this could be related to our limited number of species/replicates. Nevertheless, we observed trade-offs at the leaf and root levels between traits, namely N concentration and dry matter content. Such traits correlations between the leaf and root levels have already been reported (Freschet et al. 2010), though relatively weak relationships were found here between analogous traits belowground and aboveground. Different selective pressures for leaf and root traits as well as specialisations for the acquisition of different resources (e.g. light *vs* nutrient) could explain this absence of association between belowground and aboveground traits (Craine et al. 2005, Liu et al. 2010), while the global strategy at the plant level could remain the same since high efficiency for light or for nutrients could be related to the same physiological adaptation, as pointed out previously for stress tolerance (Chapin 1980). Although we found that leaf functional traits (LNC) can be correlated with root NH_4_^+^ and NO_3_^-^ maximal uptake rate as previously shown (Osone et al. 2008; Maire et al. 2009), here root traits (RDMC, SRL) appeared to be more related to NH_4_^+^ and NO_3_^-^ uptake rates (Rewald et al. 2014), even if deeper understanding of the relationship between root traits and nutrient acquisition remains needed (Roumet et al. 2016). The interpretation is however limited here by the fact that only three subalpine herbaceous species were studied.

### Relationship between N maximum uptake rate (Vmax) and plant strategy

Our results showed that a stronger exploitative syndrome (higher SRL, SLA, LNC and lower RDMC) was associated with higher Vmax for both inorganic N forms, rejecting the hypothesis of a trade-off between maximum uptake rate of each N forms. Ammonium toxicity has been reported for some plant species (review in Britto & Kronzucker 2002), as well as negative interactions between the uptake of NH_4_^+^ and NO_3_^-^(Kronzucker et al. 1999), and this could promote a trade-off in the acquisition of NH_4_^+^ and NO_3_^-^ between species (Maire et al. 2009). Here, we estimated NH_4_^+^ and NO_3_^-^ uptake independently to avoid such interactions during measurements, and our results did not support a trade-off but rather suggest a synergistic uptake of both N forms. Provision of NO_3_^-^ has been demonstrated to alleviate the NH_4_^+^ toxicity (Britto & Kronzucker 2002), and even to favour NH_4_^+^ uptake. We indeed observed higher uptakes for NH_4_^+^ than for NO_3_^-^, indicating a preference of all species for NH_4_^+^, especially for individuals with a more conservative syndrome of traits. This is likely to be related to the lower energetic cost for plant species to uptake and assimilate NH_4_^+^ compared to NO_3_^-^ (Salsac et al. 1987). Besides, more exploitative plants have a lower preference for NH_4_^+^compared to more conservative individuals, but expressed higher maximal uptake rates than more conservative individuals for both N forms. At the grassland plant community scale, this NH_4_^+^ *vs*. NO_3_^-^ preference is likely to have consequences on ecosystem functioning and N balance; for instance because NO_3_^-^ is more prone to leaching whereas NH_4_^+^ is better retained in soil (Boudsocq et al. 2012). Overall, our results suggest that changes in functional leaf traits related to a higher potential photosynthesis efficiency and light capture appeared to be associated at the root level with higher maximal uptake rates for both N forms.

### Nitrogen uptake variations in response to management and sampling dates

Nitrogen uptake rate is usually considered as a property of plant species, but little is known about variation in within-species N uptake rates in grasslands with different land-use history and at different times during the growing season. In our study, we observed that nitrogen uptake rates could differ strongly for the same species in different grasslands (e.g. *B. erectus* and *D. glomerata* in the UU grassland). On the other hand, the time of the year also influenced the N uptake rates of all species, with for example a higher NH_4_^+^ uptake in the autumn than in summer in UU grasslands, whereas no difference was detected in UU grasslands for *B. erectus*. Overall, grasslands were weakly discriminated by functional traits, suggesting that other factors such as soil parameters may explain the within species N uptake differences between grasslands.

Nitrogen uptake can vary depending on the amount of N available in the soil (Gavito et al. 2001). Soil NH_4_^+^ concentration, the main N source taken up by plants in our study, was similar in the four investigated grasslands, whereas a higher soil total dissolved N (TDN) was measured in the UU grassland. Consequently, the lower N uptake rates observed in this UU grassland cannot be explained by a lower N availability. As reported by previous studies, subalpine grasslands can show the legacy effects of former management activities, leading to slower N cycling (Zeller et al. 2000, Robson et al. 2007). Indeed, we observed lower pH and higher soil water and organic matter contents in the UU grassland suggesting variations in N cycling and in the quality of the available N, not only in its quantity (Garnett & Smethurst 1999, Robson et al. 2010). Supporting this hypothesis, we observed variations in soil NO_3_^-^ concentrations, and consequently soil NH_4_^+^:NO_3_^-^ ratio, between the studied grasslands. Although we could not directly relate *in situ* soil parameters to N plant uptake estimated under “controlled” conditions, we interestingly observed parallel changes for NH_4_^+^ uptake rates and soil NH_4_^+^: NO_3_^-^ ratio in grasslands where individuals have been sampled. For example, both *B. erectus* NH_4_^+^ uptake and soil NO_3_^-^ concentration were lower during the summer and higher during the autumn in TU than in UU.

Rarely investigated in natural ecosystems, experimental evidences on cultivated plants have demonstrated the effects of soil NH_4_^+^: NO_3_^-^ concentration ratio on plant N uptake (Errebhi & Wilcox 1990, Bar-Tal et al. 2001). Yet, the effects were largely species-dependent and trade off were sometime reported between NH_4_^+^ and NO_3_^-^ uptakes (Warncke & Barber 1973, Kronzucker et al. 1999, Maire et al. 2009). The preferential uptake for an inorganic N form could also be influenced by environmental and physiological factors (Britto & Kronzucker 2013). Our results did not support any trade-off in the intrinsic ability of plant species to take up both N forms, even after removing possible environmental conditions or interactions between inorganic N forms. Although we could not directly test for the relationship between soil parameters and plant N uptake rates, differences between grasslands in the N uptake within species highlight that management practices may have important effects on plant N uptake, likely through N cycling changes and the quality of the N pool available as already pointed out by previous studies (Zeller et al. 2000, Robson et al. 2007). Other studies have suggested that N preference could be dependent on the soil availability of the different N forms (Näsholm et al. 2009, Stoelken et al. 2010). While our results partially supported this hypothesis, with variation within species between different grassland, the different species sampled in the same grassland showed differences in their NH_4_^+^:NO_3_^-^ uptake ratio, supporting the hypothesis that this “preference” is partially related to the strategy of species, or at least to species identity. But overall, more exploitative species with higher maximum uptake rates for one inorganic N form are also likely to have high uptake rates for other N forms as previously found (Kastovska & Santruckova 2011).

Nevertheless, the plant preference for N forms is a complex topic (Britto & Kronzucker 2013), and careful considerations should be given to the environmental conditions where the species occur. Since N cycling is controlled by a large set of parameters including pH, soil moisture, land-use, short and long-term variations in the predominant N forms available for plants are to be expected. Under harsh conditions, plants can also take up organic N (amino acids) directly and/or through fungi (Näsholm et al. 2000). While we assumed that this source of N is of limited importance for our species in our relatively fertile grasslands (Kahmen et al. 2009), a full understanding of the N preference, and discussion about species coexistence through N forms sharing, would require careful investigations, beyond the possibility in our study. Nonetheless, the variability we observed in the ratio of uptake between the inorganic N forms suggested that, to some extent, plant physiology was adjusted to match the soil conditions where species occurred. Yet, differences between species with different strategies remain, with higher uptake rate for both N forms associated with a more exploitative strategy, and we hypothesised that this should be also the case for organic N sources (Kastovska and Santruckova 2011). Nevertheless, we acknowledge that this question could be more important in harsh environments where soil organic N is relatively more abundant as a N source for plants (Mozdzer et al 2014). Further investigations remain needed on the variations of plant N uptake under field conditions, in link with potential variations in N cycling in response to land-use or during the season (Robson et al. 2010, Legay et al. 2013).

### Variations of N uptake ability during the growing season

Plant N uptake ability also varies during the growing season, with N uptake increasing (Stahl et al. 2011) or decreasing (Jaeger et al. 1999) depending on the ecosystems investigated. In the UU grassland, NH_4_^+^ uptake was higher for all species during the autumn than during the summer, and the same was found for NO_3_^-^ uptake by *B. erectus*. Plant activity is usually considered to slow down during the autumn compared to the peak biomass in summer, an assumption supported by higher LDMC and lower SLA for all species related to the senescence of leaves. However, we did not observe any changes for root traits, suggesting that roots could remain active during this time of the growing season, especially in the process of resource storage, an important feature for subalpine/alpine plants (Kleijn et al. 2005). Additionally, studies have reported an increase of grassland N cycling rate in the autumn that could be explained by more favourable soil conditions (first rains and mild temperature), and associated with still active N uptake by plants as observed in our study (Miller et al. 2009, Larsen et al. 2012). This could also be related to the better retention of NH_4_^+^ *vs*. NO_3_^-^ in wet soils during autumn, making NH_4_^+^ more available for plant uptake (Brady and Weil 2001). Despite the fact that only few soil parameters differed between the two investigated seasons in the UU grassland, the N uptake increase in autumn was more likely a site-dependent effect related to soil conditions (Miller et al. 2009, Stahl et al. 2011, Legay et al. 2013), rather than a species response since all species did show the same pattern in the other grasslands. Yet, a multiple-year study remains necessary to better conclude on these seasonal patterns.

## Conclusion

By estimating inorganic root N uptake under controlled conditions from plants grown up under field conditions, our results support the assumption that root and leaf functional traits are associated with the ability of plants to acquire soil inorganic N. In particular, the observed pattern for roots characteristics appeared similar to the one observed in the leaf economic spectrum, with higher inorganic N uptake rates associated with more exploitative syndrome of traits. However, a weak relationship between leaf and root traits suggests that leaf traits alone were insufficient to predict inorganic N uptake. Additionally, inorganic N uptake varied within species during the growing season and in response to local conditions, making root traits and soil parameters important features of the relationships between plant functioning and grasslands N cycling. Nevertheless, these results based on excised root study need to be confirmed at the whole plant level using, for instance, ^15^N labelling.

## Data accessibility

Data are available online: http://doi.org/10.5281/zenodo.3598570

## Acknowledgements

We wish to thank Marie-Paule Bataille for IRMS analyses. The authors thank the “Conseil Régional de Basse-Normandie” for the funding of a postdoctoral position to FG. This study was conducted as part of ERA-Net BiodivERsA project VITAL, ANR-08-BDVA-008. We also thank the editor and the referees for their constructive comments which substantially improved the manuscript. Version 4 of this preprint has been peer-reviewed and recommended by Peer Community in Ecology (https://doi.org/10.24072/pci.ecology.100038).

## Conflict of interest disclosure

The authors of this preprint declare that they have no financial conflict of interest with the content of this article. François Massol and Bertrand Schatz are one of the PCI Ecology recommenders.

## Notes

http://doi.org/10.5281/zenodo.3598570

